# Ocean acidification alters the transcriptomic response in the nervous system of *Aplysia californica* during reflex behaviour

**DOI:** 10.1101/2023.11.14.567130

**Authors:** Jade M. Sourisse, Celia Schunter

## Abstract

Ocean acidification (OA) has numerous impacts on marine organisms including behaviour. While behaviours are controlled in the neuro system, its complexity makes linking behavioural impairments to environmental change difficult. Here we use a neurological model *Aplysia californica* with well-studied simple neuro system and behaviours. By exposing *Aplysia* to current day (∼500 µatm) or near-future CO_2_ conditions (∼1100 µatm), we test the effect of OA on their tail withdrawal reflex (TWR) and the underlying neuromolecular response of the pleural-pedal ganglia, responsible for the behaviour. Under OA, *Aplysia* relax tails faster due to increased sensorin-A expression, an inhibitor of mechanosensory neurons. We further investigate how OA affects habituation, which produced a “sensitization-like” behaviour and affected vesicle transport and stress response, revealing an influence of OA on neuronal and behavioural outputs associated with learning. Finally, we test whether GABA-mediated neurotransmission is involved in impaired TWR, but exposure to gabazine did not restore normal behaviour and provoked little molecular response, rejecting the involvement in TWR impairment. Instead, vesicular transport and cellular signalling link other neurotransmitter processes directly with TWR impairment. Our study shows effects of OA on neurological tissue parts that control for behaviour revealing the neurological mechanisms when faced with OA.

## Introduction

Ocean acidification (OA) can have impacts on both calcifying and non-calcifying organisms’ survival and development (1,2), physiology (3,4) and behaviour (5) observed in several marine taxa (6). Fish exhibit behavioural impairments amongst others in antipredator response (7–9) and learning (10,11). However, invertebrates such as snails and crabs have also displayed behavioural impairments caused by OA (12–14). Given that behaviour can significantly impact population dynamics (15–17), it is crucial to understand the drivers and underlying mechanisms to predict the trajectories of behavioural responses in a rapidly changing environment (18,19).

The alteration of behaviour is associated with the brain or neuronal cells with a change occurring to the molecular drivers (such as gene expression). One proposed cause of behavioural impairments in a reduced pH environment is altered GABA neurotransmission (20). GABA is a major inhibitory neurotransmitter found across the animal kingdom (21–23) and GABA_A_ receptors are ion channels whose activation in normal conditions results in neuron inhibition. When the partial pressure of CO_2_ (pCO_2_) is elevated in seawater, the acidosis response of marine animals modifies internal ion gradients and the subsequent activation of GABA_A_ receptors results in neuron excitation (20). This hypothesis was built on the fact that animals reared in elevated CO_2_ conditions had behavioural impairments at least partially restored when treated with the GABA_A_ receptor antagonist gabazine (13,24,25). Changes in the expression of genes involved in GABAergic neurotransmission in response to OA have also been found (26–29), supporting that elevated pCO_2_ acts in neuronal cells at the molecular level, affecting pathways throughout the brain which in turn can produce impairments at the whole organism level. Nevertheless, the complexity of the vertebrate brain brings difficulties in understanding how molecular alterations inside neurons caused by environmental changes such as OA modify the course of information transmission across synapses and throughout regions of the nervous system, *in fine* driving behavioural changes.

One way of linking environmental influence with nervous system functioning and its behavioural output is through the study of a simple and well-studied neuro system, such as that of the California sea hare (*Aplysia californica*). Contrary to vertebrate brains, the nervous system of *Aplysia* is composed of only 20,000 large neurons (30). Several of its behaviours have been characterized in neural networks that link the nervous system and sensory organs to muscles (31,32). As a result, it is possible to obtain the gene expression profile of specific nerve ganglia which control for specific behaviours. In addition, *Aplysia* displays a similar “acid–base regulator” profile to that of fishes during OA by accumulating bicarbonate (HCO_3_^-^) ions (33). Since invertebrate GABA also binds to HCO_3_^-^ permeable ionotropic GABA receptors (34), the hypothesis of OA-altered GABA neurotransmission causing behavioural impairments should be applicable in this model species. One suitable behaviour performed by *Aplysia* to test this hypothesis is the Tail Withdrawal Reflex (TWR): during the TWR, the tail is withdrawn for protection after a tactile stimulus is applied to it (35,36). It is a behaviour for which controlling neurons in the pleural-pedal ganglia have been extensively described (35,37,38), with neuron populations sensitive to several specific neurotransmitters such as glutamate (39), dopamine, FMRFamide (40) and GABA (41). The TWR was also shown to be impaired by OA itself (33). Finally, TWR can be involved in non-associative learning, notably reflex habituation (42), which is likely caused by a presynaptic mechanism preventing transmitters release (43).

We investigate the molecular processes affected by OA in the nervous system of the California sea hare (*Aplysia californica*) as it performs a “simple” behaviour. We observed the TWR response when animals were reared in either control (∼500 µatm) or predicted near-future (∼1100 µatm) CO_2_ conditions. This further allows us to investigate whether OA influences learning experiences and the involvement of GABAergic neurotransmission in behavioural impairments. Our three experiments focused on (1) how the innate TWR behaviour changes with elevated CO_2_ and the underlying mechanisms in the nervous system are, (2) whether habituation of the TWR is influenced by elevated CO_2_ and what corresponding neuromolecular changes take place in the reflex’s circuitry and (3) the potential role of GABAergic neurotransmission in the OA-induced behavioural changes of *Aplysia* (Fig. 1). Together, these experiments pinpoint the molecular basis driving behavioural modifications when faced with future ocean acidification.

**Figure 1:**
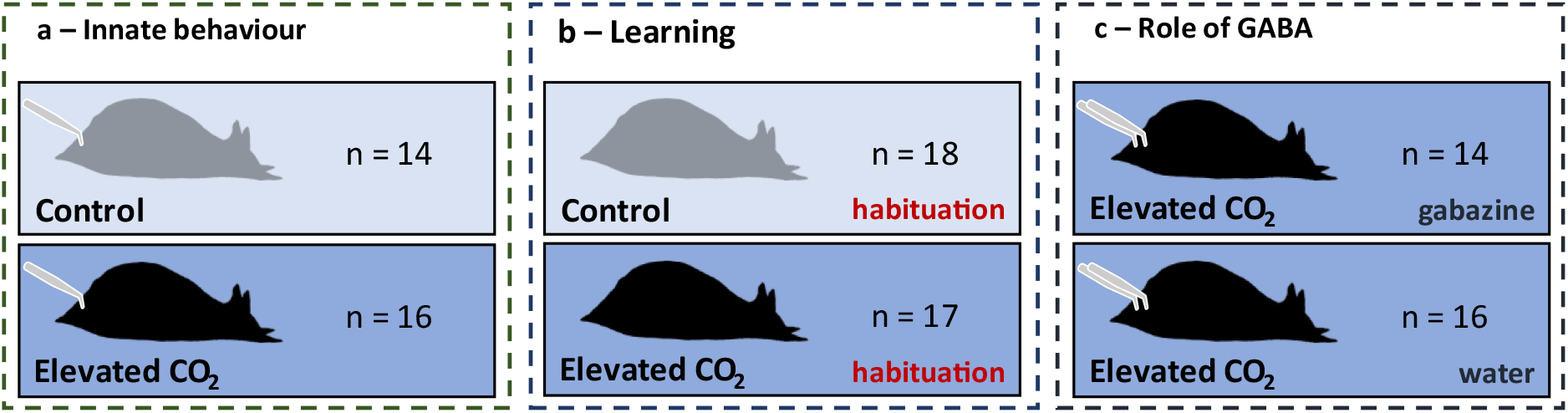
Experimental design for the study of Aplysia’s Tail Withdrawal Reflex (TWR) response under elevated CO_2_ conditions. The TWR is elicited by a tap on the tail with a needle. In the learning experiment (b), animals received a habituation training and TWR responses were observed pre- and post-training. In the GABA experiment (c), animals were either momentarily exposed to gabazine or to control water. The number of biological replicates is indicated by the n value for each group.

## Methods

### Experimental design

Adult sea hares weighing 52.8 ± 9.8g were imported from the National Resource for Aplysia (University of Miami, USA), and acclimated to their new environment for three days in ambient control conditions. After this acclimation period they were exposed to either control (pCO_2_ ∼ 450 µatm; n = 36) or near-future CO_2_ conditions (pCO_2_ ∼ 1200 µatm; n = 54) for seven to ten days at the University of Hong Kong. The elevated CO_2_ level was chosen following the RCP 8.5 scenario of the IPCC (44). Animals were housed in flow-through systems with natural seawater from Hong Kong Island shores (Fig. S1). Exposure of animals to elevated CO_2_ was created through bubbling a mix of air and CO_2_ gases at a rate of 15L/min, controlled by a PEGAS 4000 MF Gas Mixer (Columbus Instruments). To reach the desired pCO_2_, the mixing parameters were set as follows: air flow was at 15 000 cc/min, CO_2_ flow (vCO_2_ in cc/min) was calculated as a function of the CO_2_ desired partial pressure (pCO_2_ in %), the CO_2_ ambient partial pressure was at 0.04 % and the gas mix flow rate was 15L/min. The exposure to elevated CO_2_ was gradual and consisted in a first exposure to 700 µatm for two days, then 1000 µatm for four days, to then finally reach the target value of 1200 µatm for seven to ten days (depending on the experiment, see next section). The pH was measured daily in randomly chosen tanks from both control and treatment groups using a WP-91 waterproof pH meter (TPS) and a Seven2GO pH meter (Mettler Toledo) and the total alkalinity (TA) was measured weekly using a G20S titrator (Mettler Toledo). Using the titrator’s measurements, pCO_2_ values were calculated using the CO2_SYS_ software (45). Following the recommendations of the National Resource for *Aplysia*, temperature was set at 16°C using HC-1000A chillers (Hailea) and measured daily in randomly chosen tanks from both control and treatment groups using a WP-91 waterproof thermometer (TPS) and a Seven2GO thermometer (Mettler Tolder). Salinity was monitored daily using a portable refractometer. Weekly measurements of nitrate in randomly chosen tanks from both control and treatment groups were done using a HI97728 nitrate photometer (Hanna Instruments) to ensure good water quality. The animals were kept under a 12/12h light-dark cycle. *Aplysia* were fed *Agardhiella subulata* algae every 3 days as instructed by the National Resource for *Aplysia*.

Across all tanks, the salinity was at 32 ± 1.6 ‰ and the temperature was at 15.5 ± 0.6°C for the whole experimental duration (Suppl. Table 1). The elevated pCO_2_ conditions over the final seven to ten days of experiment were 1100.3 ± 59.9 µatm, whereas the control pCO_2_ was 497.5 ± 57.4 µatm (Fig. S2). pCO_2_ was significantly different between treatment groups (Wilcoxon rank sum test, p-value < 0.001). Additionally, there was no effect of the rearing system identity on pCO_2_ within the “treatment” group (ANOVA, p-value = 0.628).

### Behavioural assays

All behavioural assays were performed by video recording using a Canon EOS M50 camera. To test the innate reflex response to Tail Withdrawal Reflex (TWR) after elevated CO_2_ exposure of seven days, animals reared either in control (n = 14) or elevated CO_2_ conditions (pCO_2_ ∼ 1200 µatm; n = 16) were acclimated for 30 min in the behavioural experimental tank (Fig. 1a). The TWR was then elicited three times per animal, by pressing onto the tail with an 20G gauge needle and with each assay separated by ten minutes (Fig. 1a), as performed in previous studies investigating the TWR (33,36).

To test if learning is impaired under OA conditions, *Aplysia* either reared in control (n = 18) or elevated CO_2_ conditions (n = 17) received a habituation training after a ten days exposure in their rearing tanks (Fig. 1b). A period of 30 min for acclimation to the experimental setup was given to individuals, and then the pre-training response was assessed three times the same way as for the innate experiment, with a resting period of five min each. Training was then provided by delivering 30 repeated tail stimuli, with 30s intervals between each. The post-training TWR response was assessed 20, 25 and 30 min after training in the same way as pre-training. This protocol is the same as that of another previous study investigating habituation of the tail withdrawal reflex (46).

To test if GABA-ergic neurotransmission is involved in observed behavioural impairments, *Aplysia* reared under near-future CO_2_ conditions (pCO_2_ ∼ 1200 µatm) for seven days were either administered the GABA_A_ receptor antagonist gabazine (4 mg/L of SR-95531, Sigma; n = 14), or control water solution (n = 16; Fig. 1c). The administration route was dissolution through the experimental tank water. The TWR behaviour was recorded after a 30 min acclimation period similarly to the innate experiment.

The reflex response was measured from video recordings in terms of duration by an observer blind to experimental conditions (Table S2). Measurements were discarded either if the tail was not clearly visible after retraction, or if the TWR triggered inking, or if the reflex was a Siphon Withdrawal Reflex. Mean durations were compared between control and treatment groups for each experiment. All statistical were performed in R (R Core Team, 2018).

### RNA sequencing and gene expression analysis

All individuals tested served for collection of RNA from the pleural-pedal ganglia, the nervous tissue part known to be in control of the TWR’s circuitry with sensory neurons located in the pleural ganglia and motoneurons found in the pedal ganglia (31,47–49). The animals were dissected immediately following their behavioural assay and for each animal the pleural-pedal ganglia were snap frozen in liquid nitrogen and kept at -80°C until RNA extraction. The pedal-pleural ganglia were then homogenized using sterile silicon beads for one minute at highest speed in a Tissue Lyzer (Qiagen) and RNA was extracted following the TRIzol™ Plus RNA Purification Kit (Thermo Fisher). The RNA concentration and quality of some samples was measured using TapeStation (Agilent) to ensure good sample quality. Samples were sequenced at 150bp paired end on an Illumina NovaSeq at the Centre for PanorOmic Sciences (CPOS) of the University of Hong Kong. After sample sequencing, raw sequence data (on average 38.5± 3.9M reads per sample; Table S3) were trimmed off adapters and filtered based on read quality using Trimmomatic (50). Trimmomatic was ran using the following parameters: “ILLUMINACLIP: all_PE.fa:2:30:10:8:TRUE LEADING:4 TRAILING:3 SLIDINGWINDOW:4:20 MINLEN:30”. Then, the filtered data (on average 36.5± 3.7M reads per sample; Table S3) was mapped against the reference genome AplCal3.0 and its RefSeq annotation available for *Aplysia* (51), using the program HISAT2 (52).

Differential expression (DE) analyses were led using DESeq2 v 1.38 (53) to investigate which genes were differentially expressed between *Aplysia* reared either in control or elevated pCO_2_. A Likelihood Ratio Test (LRT) was used to ensure that neither the system nor the tank identity were factors of influence on gene differential expression. Then, a Wald test was performed to identify DE genes depending on the factor of interest. The design formula of innate and learning experiments (“∼ pCO_2_”) allowed pairwise comparisons of groups depending on the two pCO_2_ levels, whereas the design formula of the GABA experiment (“∼ Gabazine”) allowed pairwise comparisons between *Aplysia* either exposed to gabazine or control water, all at elevated pCO_2_. Differentially expressed genes with a baseMean under 10 and/or an absolute value of log2foldchange inferior to 0.3 were discarded to ensure that differential expression was not an artifact of low counts, and to increase stringency. Additionally, to identify which genes have their expression patterns possibly mediated by either the TWR response, CO_2_ and/or habituation, and could in turn play a role in the behavioural change of *Aplysia*, a weighted gene co-expression network analysis (WGCNA) was performed using the WGCNA v 1.72.1 package (54). Mean TWR duration of each individual and final target pCO_2_ were provided as trait data, as well as binary encoded information regarding their habituation status (“0” naïve, “1” habituated) and gabazine exposure (“0” not exposed to gabazine; “1” exposed to gabazine). The following parameters were used to build the network: power = 7 (with R^2^ > 0.90), TOMType = “signed”, minModuleSize = 30, reassignThreshold = 0, mergeCutHeight = 0.25, verbose = 3. Clusters of genes whose expression patterns were correlated with either pCO_2_ and/or habituation along with TWR duration, or with TWR duration alone were identified. For significant differentially expressed (DE) genes and for genes highlighted by the WGCNA analysis, functional enrichment analyses were performed using OmicsBox v (Fisher’s Exact Test). The GO annotations used for the enrichment analysis were retrieved from BioMart in OmicsBox, using the “Database of genes from NCBI RefSeq genomes IDs”. The Gene Ontology (GO) terms with an FDR adjusted p-value below the 0.05 threshold were considered enriched among the DE genes, and the list of GO terms was reduced to its most specific. Uncharacterized loci highlighted by the gene expression analyses were searched in Rapid Ensembl release 109 (55) in the *Aplysia californica* genome AplCal3.0 (assembly GCA_000002075.2) to predict their putative function.

## Results

### Behavioural response to ocean acidification during TWR

In the innate experiment, *Aplysia* exposed to elevated CO_2_ conditions relaxed their tail after withdrawal 29% faster with 12 ± 5 (mean ± SD) seconds to relax, which was significantly different from control condition which relaxed after 17 ± 6 seconds (Wilcoxon rank sum test, p-value < 0.001; Figure 2a). In the habituation experiment, we saw the same significant effect of CO_2_ exposure on Tail Withdrawal Reflex (TWR) duration before habituation (Wilcoxon rank sum test, p-value = 0.0216; Fig. S4), as *Aplysia* exposed to elevated pCO_2_ before conditioning took on average 12 ± 5 seconds to relax in elevated CO_2_ but 15 ± 7 seconds in control condition.

**Figure 2:**
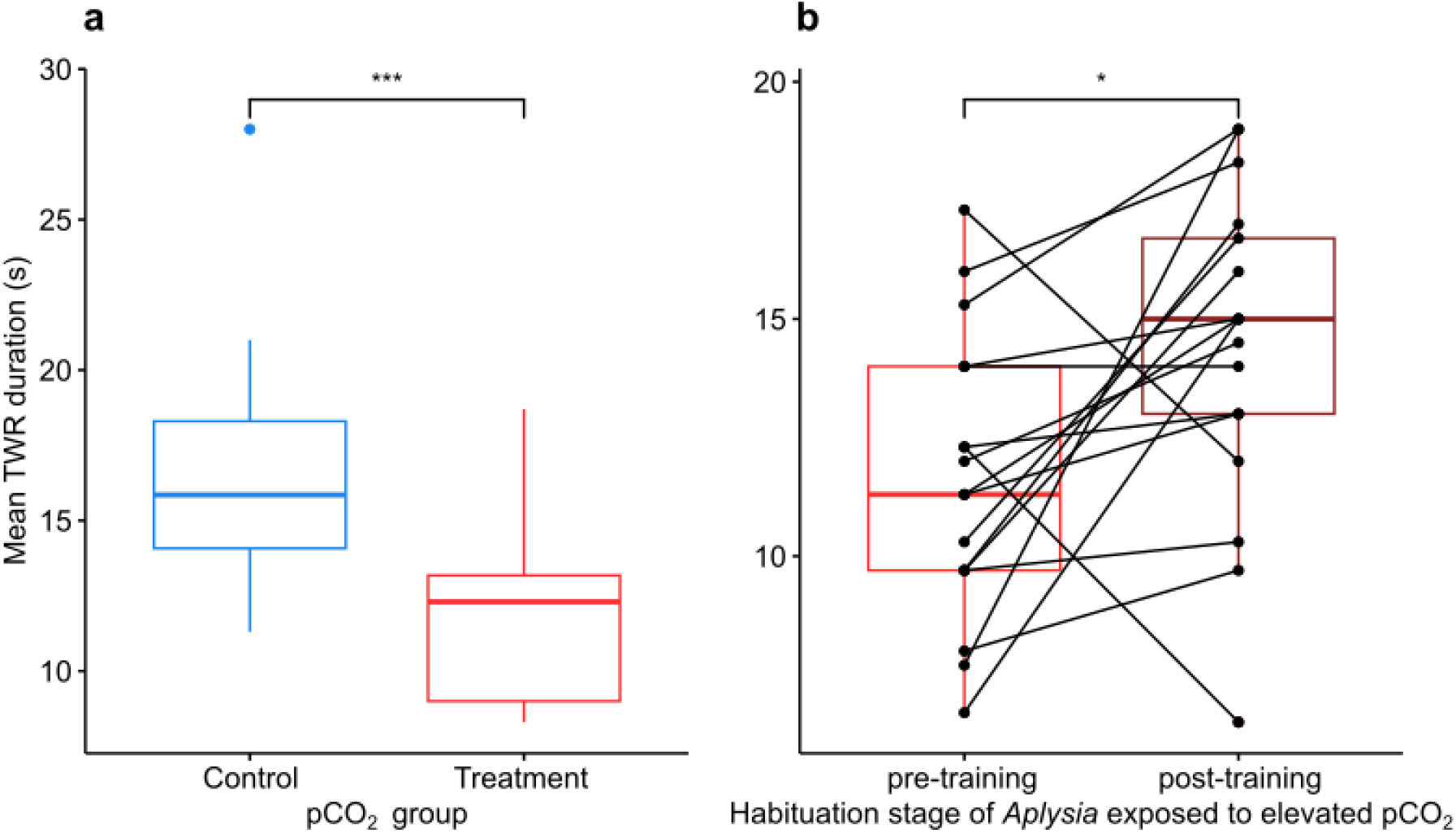
Tail Withdrawal Reflex (TWR) duration (s) of Aplysia in the innate experiment (a) as a function of their pCO_2_ conditions (control ∼500 µatm or treatment ∼1100 µatm) and in the habituation experiment (b) reared at elevated pCO_2_, before (pre-training) and after (post-training) habituation training; stars (*** or *) indicate the level of significant difference between the mean values

Habituation caused *Aplysia* to relax their tail in 14 ± 6 s for both control and elevated pCO_2_ conditions. While it was not significantly different to pre-training for control individuals (Wilcoxon rank sum test, p-value = 0.467; Fig. S5), habituation surprisingly increased TWR duration for all but two individuals exposed to elevated CO_2_ (p-value = 0.0238; Fig. 2b). Elevated CO_2_ caused the tail to relax approximately 17% slower after conditioning. Finally, in the GABA experiment where all individuals were exposed to elevated CO_2_, there was no significant effect of gabazine on TWR duration (Wilcoxon rank sum test, p-value = 0.1857; Fig. S3). Individuals who were administered control seawater took on average 12 ± 5 seconds to relax from TWR whereas individuals who were administered gabazine relaxed after 14 ± 6 seconds.

### Molecular response to ocean acidification during TWR

Acidification provoked a large transcriptomic response in the TWR-mediating parts of the *Aplysia* nervous system, with a total of 1761 genes significantly differentially expressed between naive *Aplysia* exposed to different pCO_2_ levels, while 725 genes were differentially expressed among habituated *Aplysia*. Of these, 248 genes were commonly differentially expressed due to acidification. This general response to ocean acidification affected calcium ion (Ca^2+^) binding (Tables S4 & 5), through differential expression of receptors of Ca^2+^ important to nervous system functioning. Calcium transport exhibited downregulation of calcyphosin-like gene possibly regulating ion transport and a trimeric intracellular cation channel gene (*tric-1B.2*), which facilitates the active intracellular release of Ca^2+^ (Tables S6 & 7). Additionally, calmodulins, which are calcium-binding messengers involved in calcium signal transduction and synaptic plasticity (Tables S6 & 7), and calcium-binding protocadherins involved in cell-cell interactions were also differentially expressed (Tables S6 & 7). Related to calcium signalling, ocean acidification also provoked changes in excitatory neurotransmission with the upregulation of neuronal acetylcholine (ACh) receptor subunits, the Cerebral Peptide 2 precursor (*CP2PP*), the neuropeptide 15 receptor (*npr-15*), the neuropeptide CCHamide-1 receptor and an ionotropic glutamate receptor (Fig. 4a, Table S5). Furthermore, pCO_2_ levels positively correlated with an ACh receptor subunit precursor (Table S9). Another excitatory ionotropic glutamate receptor of the AMPA type was predicted for an uncharacterized locus (*LOC101845288*) in the genome which had the highest expression level among all transcripts yet was significantly downregulated due to acidification (Fig. 3b and 4a, Tables S6 & 7) in both naïve and habituated *Aplysia*. Acidification specifically had an effect on serotonergic neurotransmission in naïve *Aplysia* as the expression of two serotonin receptors with excitatory activity on neurons (*5-HT_2_* and 5-HT_4_) but also one inhibitory serotonin receptor (5-HT_1D_), was expressed at higher levels at elevated pCO_2_ (Fig.4b, Tables S5 & S9). Another gene involved in inhibitory neurotransmission was the upregulation of BTB/POZ domain-containing protein *KCTD12*, which is a GABA_B_ receptor subunit (Table S6). Finally, acidification caused the upregulation of a calcium-binding inhibitory co-transmitter released by mechanosensory neurons, the precursor mRNA of sensorin-A (*psc1*; Figure 3a & 4a; Table S6).

**Figure 3:**
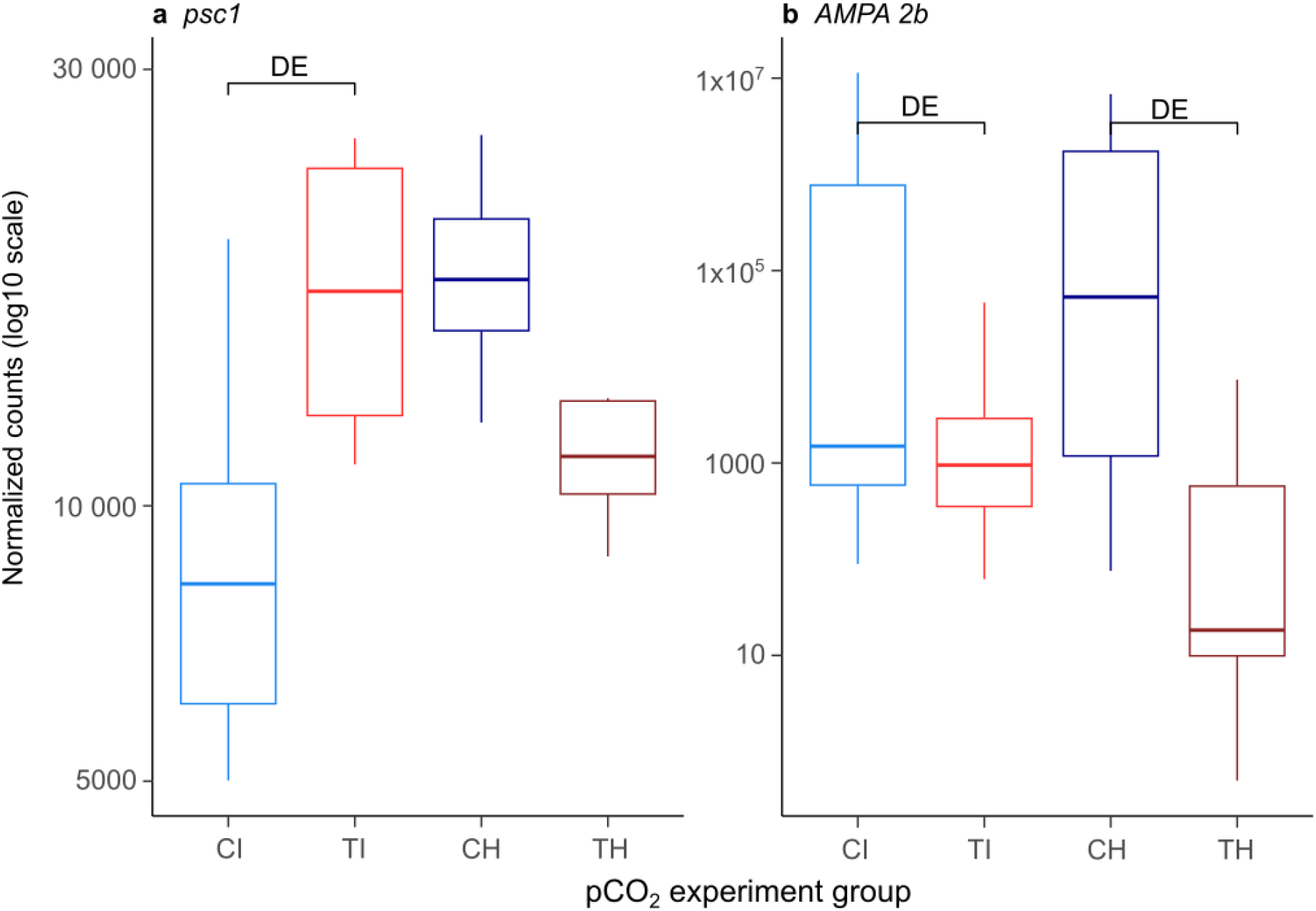
Normalized counts (log_10_) in all pCO_2_-experiment groups for the gene psc1 (a) and the AMPA 2b predicted coding gene (b); Aplysia were grouped as a function of the pCO_2_ they were reared at and the experiment they participated in: CI = Control (∼500 µatm) Innate, TI = Treatment (∼1100 µatm) Innate, CH = Control Habituated, TH = Treatment Habituated; “DE” shows which pairwise comparison revealed the gene as differentially expressed

**Figure 4:**
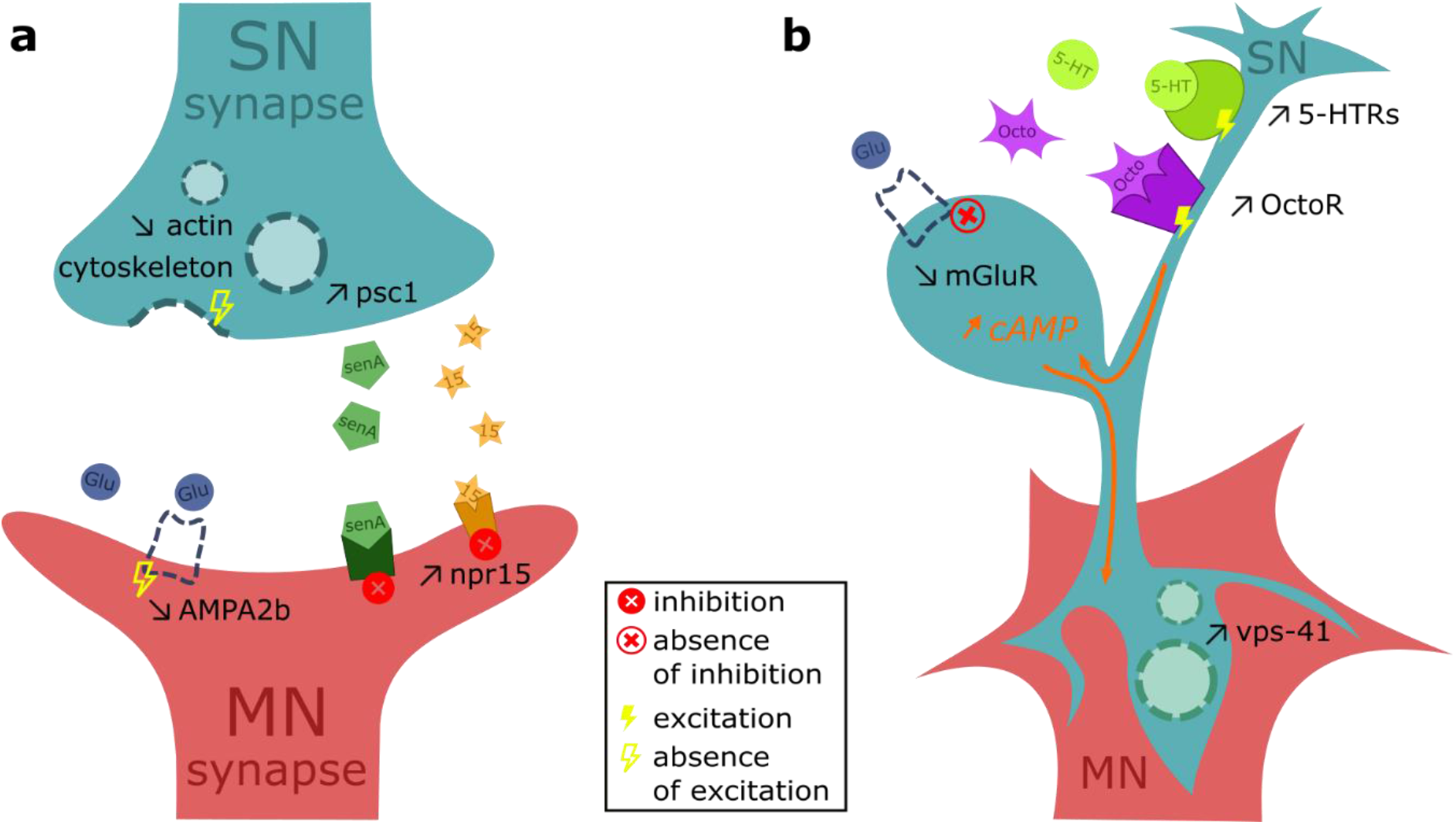
Molecular mechanisms in the TWR neural circuitry of Aplysia when exposed to elevated pCO_2_ (a) during innate elicitation of the reflex and (b) after habituation of the reflex. SN = sensory neuron; MN = motoneuron; npr15 = neuropeptide 15 receptor; psc1 = pleural sensory cluster 1 gene, precursor of senA = sensorin A; Glu = glutamate; AMPA2b = AMPA 2b receptor gene; 5-HT = serotonin; 5-HTR = serotonin receptor; Octo = octopamine; OctoR = octopamine receptor; mGluR = metabotropic glutamate receptor; cAMP = cyclic adenosine monophosphate; vps-41 = vacuolar protein sorting protein 41 gene

Ocean acidification also affected components of the cytoskeletal network. Notably, the expression of actin-related genes was significantly positively correlated with mean TWR duration and negatively correlated with pCO_2_ (Fig. S6; Table S8), including Actin related protein (ARP) 2/3 complex subunits, ARPs, an F-actin-capping protein subunit, a myosin-10, a FLII actin remodelling protein, a plastin-1 and one actin protein (Table S9). Furthermore, genes involved in the actin-myosin interaction regulating cell morphology and cytoskeletal organization were differentially expressed, such as myosin light chains coding genes (Tables S6 & 7). Specifically in naïve *Aplysia*, an actin-interacting protein, ARP 2/3 complex subunits and the PRKCA-binding protein (*pick1*) coding genes, which interact to regulate excitatory synaptic plasticity, were downregulated (Table S6). Other genes associated with cytoskeletal motion and intracellular transport were downregulated at elevated pCO_2_, such as genes coding for tubulin chains as well as the centrin-3 and EF-hand domain-containing protein coding genes which are involved in microtubule organization (Tables S6 & 7). Finally, genes interacting with dynein also involved in cytoskeletal motion were consistently affected by acidification, such as a parkin coregulated gene homolog, a dynein assembly factor, dynein chains and dynein regulatory complex proteins (Tables S6 & 7).

Several energy related processes were affected by acidification. Firstly, ATP metabolism through proton transmembrane transport, proton-transporting ATPase activity and acyltransferase activity (Table S4) were altered in naïve *Aplysia* with the downregulation of ATP synthase, ATPase, and ATP-citrate synthase and citrate synthase coding genes (Table S6). Biosynthesis of nucleic acids, which is energetically expensive, was suppressed (Table S4) and protein metabolism was affected by acidification through the downregulation of all genes involved in alpha-amino acid synthesis (Tables S4, 6 & 7) tRNA ligases, initiation factor subunits and elongation factors (Tables S6 & 7) in both naïve and habituated *Aplysia*. Protein synthesis itself was altered through the differential expression of ribosomal protein coding genes (Table S6) as well as the post-translational modification glycosylation due to downregulation of protein glycosyltransferase subunits (Tables S6 & 7). Finally, downregulation of protein disulfide-isomerase coding genes (Table S6) impacted unfolded protein binding in naïve *Aplysia* exposed to elevated pCO_2_ (Table S4).

Acidification triggered changes in the expression of cellular stress response genes. Heat shock proteins 70 (B2 subfamily) were upregulated in *Aplysia* exposed to elevated pCO_2_ (Tables S6 & 7) and genes with antioxidant activity (Table S4) were downregulated such as glutathione S-transferases, a glutathione reductase, a carbonyl reductase [NADPH] 1 isoforms, glutaredoxin and peroxiredoxins (Tables S6 & 7).

Habituation training revealed on effect of acidification on learning experiences in the nervous system of *Aplysia*. After *Aplysia* received habituation training, exposure to elevated pCO_2_ did not alter cell signalling and organization functions as seen in untrained *Aplysia*. Interestingly, the precursor mRNA of sensorin-A (*psc1*) was upregulated due to acidification in naïve *Aplysia,* but habituated *Aplysia* had elevated levels of this gene no matter the pCO_2_ (Figure 3a & 4a; Table S6), suggesting that overexpression of *psc1* could also be achieved through habituation training. Similarly, only in naïve *Aplysia*, acidification provoked changes in the expression of genes involved in cell signalling processes mediated by GTP binding and GTPase activity (Table S4), with downregulation of G-proteins involved in vesicular traffic such as ADP ribosylation factor (ARF), ARF-binding protein (*GGA1*) and two ADP-ribosylation factor-like (ARL) proteins (Table S6), which were not altered by OA in habituated *Aplysia*. Further processes seen for naïve *Aplysia* with OA, but not in habituated *Aplysia* despite the elevated CO_2_ levels were intracellular transport, notably vesicle-mediated and protein transport (Fig.4a, Table S4). Expression in genes coding for vacuolar protein sorting-associated proteins, YIPF proteins, coatomer subunits and exocyst complex components, as well as a vesicle-associated membrane protein (VAMP)/synaptobrevin-binding protein, a synaptobrevin and a synaptobrevin homolog, downregulated for naïve *Aplysia*, are at control levels in habituated *Aplysia* (Table S6). Finally, otoferlin, involved in the Ca^2+^-triggered synaptic vesicle-plasma membrane fusion and in the control of neurotransmitter release at output synapses, was upregulated only in naïve *Aplysia* but not in habituated OA *Aplysia* (Table S6). Further evidence that habituation and pCO_2_ affect cell signalling differently was found as co-expressed genes positively correlated with pCO_2_ but negatively correlated with habituation training participated in GTPase mediated signalling (Table S8) such as four guanine nucleotide exchange factor coding genes and two genes coding for dedicators of cytokinesis (Table S9). Furthermore, genes of the thioredoxin antioxidant system and further antioxidant enzymes (glutathione peroxidase and superoxide dismutase) were at control levels for habituated *Aplysia* but downregulated in naïve *Aplysia* (Table S6) and these genes were also positively correlated with mean TWR duration and habituation, but negatively correlated with pCO_2_ (Fig. S6; Table S9). Overall, genes involved in cell signalling and cellular stress response whose expression were altered by acidification in naïve *Aplysia* were not differentially expressed in habituated *Aplysia*, indicating a possible “cancelling/antagonistic” effect of habituation training on gene expression under acidification.

Habituation training, however, also caused specific effects associated with acidification in the nervous system of *Aplysia*. Signalling and vesicle transport were affected by the pCO_2_ level, but the genes involved in these functions were different after they experienced habituation training. For instance, regucalcin, which is a suppressor protein of cell signalling, was downregulated by acidification only in trained *Aplysia* (Table S7). Moreover, five genes involved in cellular transport were upregulated with elevated pCO_2_ only in habituated *Aplysia*, notably two *vps-41* isoforms, one VAMP coding gene, one *VTI1B* homolog and one gene coding for an ARL protein 6-interacting protein (Fig. 4b, Table S7). GTP-mediated signal transduction was also differently affected by pCO_2_ after habituation, as the small GTPase *Rap*, a Ras-related protein (*Rab-21*) involved in membrane trafficking, the *IQGAP1* gene regulating the dynamics of the actin cytoskeleton and the Ras Guanine nucleotide exchange factor (GEF) 1B were all upregulated in trained *Aplysia* (Table S7). Furthermore, genes involved in neuromodulation were downregulated at elevated pCO_2_ only in habituated *Aplysia*, such as two metabotropic glutamate receptors involved in the regulation of neurotransmitter release notably by preventing cAMP production (Fig. 4b, Table S7). Further cAMP-responsive pathway genes including CREB-regulated transcription coactivator and CREB3 regulatory factor were upregulated specifically after habituation training (Table S7) and the CREB3 regulatory factor was also positively correlated with both pCO_2_ and habituation, alongside the cAMP response element-binding protein (CREB) coding gene (Table S9). Both pCO_2_ levels and habituation training positively also correlated with the expression of serotonin and octopamine receptors involved in neuromodulation (Fig. 4b, Table S9). Finally, genes involved in oxidoreduction and cellular stress response were differentially expressed depending on pCO_2_ only in habituated *Aplysia* such as the cytochrome b, the cytochrome c oxidase and nitric oxide synthase coding genes (Table S7). Therefore, habituation training triggers molecular changes specifically when individuals are experiencing acidification, such as modified expression of genes performing cell signalling, neuromodulation and oxidoreduction functions.

The potential role of GABA-ergic neurotransmission in the observed behavioural impairments due to acidification was investigated by administering gabazine to *Aplysia* reared under near-future CO_2_ conditions. Only 20 genes were differentially expressed due to the administration of gabazine in comparison to control water (Table S10). All except one (uncharacterized) were upregulated in *Aplysia* exposed to gabazine. They coded for hydrolases, proteases, kinases, a junction-mediating and regulatory protein (JMY), a cytochrome P450 2D26-like protein and a betaine--homocysteine S-methyltransferase 1 (Table S10).

## Discussion

Our study connected the behavioural impairment caused by OA in *Aplysia californica* with the molecular changes occurring in the relevant part in the nervous system and thereby pinpointing the underlying mechanisms of OA-driven behavioural alterations. Acidification significantly decreased Tail Withdrawal Reflex (TWR) duration consistent with a previous study (33) and provoked large changes in gene expression inside the TWR mediating parts of the nervous system, the pleural-pedal ganglia. Several of those observed changes constitute possible molecular mechanisms causing the behavioural impairment, such as the upregulation of the peptide sensorin-A (*psc1*) coding gene. As *psc1* is an inhibitory transmitter in *Aplysia* mechanosensory neurons (56), CO_2_-mediated increases in sensorin-A mRNA expression can cause downstream neurons of the circuit to be inhibited, therefore not transmitting the stimulus information to the end of the reflex circuitry which is responsible for prolonged contraction. Hence, by leveraging the intricate knowledge of the TWR circuit in the simple neuro system of *Aplysia* we are able to connect the behavioural impairment to underlying mechanisms.

Further neuronal signalling inhibition may be at play, however, as we find a neuropeptide receptor (*npr-15*) and an AMPA receptor changed expression levels when exposed to elevated CO_2_. These alterations could lead to an increase in inhibitory response, such as seen in an octopaminergic manner for *C. elegans* in the case of *npr-15* (57) with increased inhibition of neurons inside the reflex circuitry. Furthermore, reduced glutamatergic excitatory transmission via downregulation of the AMPA receptor may lead to decreased neuron excitability in *Aplysia* (36,58) resulting in less firing of motoneurons for prolonged foot muscle contraction. Differential expression of receptors involved in neurotransmission can therefore lead to the decreased TWR duration caused by acidification.

Apart from neurotransmission, cellular organisation and signalling are altered by elevated CO_2_ exposure and could be involved in the decreased TWR duration, notably genes involved in actin dynamics were correlated with TWR duration. The actin cytoskeleton regulates synaptic areas morphology and synaptic vesicle pools (59,60) and it is sensitive to ocean acidification in other molluscs (61,62). Synaptic vesicle mobilization in particular was found to be sensitive to acidification in nerve terminals of vertebrate motoneurons (63), as an increase in intracellular proton concentration suppresses synaptic vesicle delivery. In the case of *Aplysia* and fishes, even though acid-base regulation occurs following exposure to elevated CO_2_, maintaining acid-base balance is achieved through retention and/or uptake of HCO_3_^-^ to maintain the internal pH despite elevated proton concentrations (33). Since internal proton concentration is maintained at a high level, synaptic vesicle delivery could be suppressed through the observed reduced expression of actin-related genes involved in the mobilisation of synaptic vesicles. As a result, this reduced vesicle activity would *in fine* inhibit neurotransmission and decrease the reflex response and duration. Furthermore, other cytoskeletal motors, such as tubulin, were also downregulated by elevated pCO_2_. This same pattern has been found in tubeworm and oyster larvae (64,65). Evidence of changes in cytoskeleton components during exposure to elevated pCO_2_ suggests that acidification not only interferes with normal vesicle functioning at the synapse, causing behavioural impairment, but might also alter numerous other functions performed by the cytoskeleton. For example, in blood clams changes in the cytoskeleton due to acidification may have implications on the immune response because cytoskeletal components perform phagocytosis (66). Hence, cytoskeletal changes may reduce synaptic mobilization which is a further mechanism for TWR changes under acidification conditions, but modified cytoskeleton organization could also cause other downstream effects at the whole organism level.

The decreased synaptic activity among neurons of the reflex circuitry with acidification may have led to compensatory mechanisms through differential expression of genes involved in neurotransmission. One potential compensation is the upregulation of acetylcholine receptors in *Aplysia* pedal-pleural ganglia, also observed in pteropod nervous systems under ocean acidification (67,68). The upregulation of neuronal acetylcholine could increase Ca^2+^ levels and intensify calcium-mediated exocytosis of synaptic vesicles, which can be an alternative way of restoring neurotransmitter delivery impaired by the downregulation of genes involved in synaptic mobilization in *Aplysia*. Additionally, with otoferlin placing synaptic vesicles near calcium channels to allow fast exocytosis of neurotransmitters (69), the observed upregulation of otoferlin in *Aplysia* at elevated pCO_2_ could also facilitate Ca^2+^-mediated release of synaptic vesicles from the sensory neurons and restore a flow of neurotransmitters in the synaptic area. Furthermore, the upregulation of genes involved in the recycling of synaptic vesicles could be a compensatory mechanism in *Aplysia* when faced with acidification. Maintenance of neurotransmission by recycling existing receptors could counteract decreased gene expression of such receptors. For example, the upregulation of the huntingtin-interacting protein and AP180 coding genes in *Aplysia* neurons at elevated pCO_2_ could maintain neurotransmission by enhancing the recycling of already existing AMPA receptors (70,71), despite the downregulation of the AMPA receptor coding gene that would limit its neobiosynthesis. Finally, increased production of neuropeptides, such as those derived from the Cerebral Peptide 2 precursor (*CP2PP*) could also be a way to compensate for impaired neurotransmission. *CP2PP* is a precursor to ten bioactive neuropeptides in *Aplysia*, among which some act on the foot muscle (72,73), therefore its upregulation could lead to increased production of neuropeptides which could activate tail motoneurons. Overall, *Aplysia* shows signs of transcriptional reprogramming to compensate for reduced neurotransmission when exposed to acidification in the nervous system.

The molecular response to acidification in the pedal-pleural ganglia also comprised several important functions such as ATP, nucleic acid and protein metabolism. Downregulation of metabolism genes in *Aplysia* could be a sign of metabolic suppression as commonly seen in other invertebrates when exposed to elevated pCO_2_ (74–77), implying that downregulation of oxidative metabolism genes in sensitive organisms might compromise the cellular stress response (78). However, in *Aplysia*, the downregulation of oxidative metabolism genes under OA is not sufficient to imply that the nervous system may be more susceptible to cellular stress, because the upregulation of heat shock protein genes 70 was also observed. This type of heat shock protein is expressed when cellular protection is required, notably but not restricted to oxidative stress conditions (79). Their upregulation in *Aplysia* to protect cells against OA-mediated stress has also been documented in Sydney rock oysters (80). Hence, instead of metabolic suppression, the metabolic downregulation may rather point to reallocation of energy towards homeostasis. This has been suggested as a cellular strategy to redirect the energy in the most effective way possible towards immediate essential processes at the expense of other functions (77). *Aplysia* exposed to 1200 µatm of pCO_2_ are able to maintain their haemolymph pH_e_ levels similar to that of control by accumulating HCO_3_^-^ (33). It is therefore possible that the downregulation of genes involved in expensive processes is a means to ensure that sufficient resources are allocated to the uptake of HCO_3_^-^ inside the haemolymph to maintain acid-base homeostasis at the whole organism level.

When *Aplysia* were trained under OA, contrary to our expectations, the TWR duration increased. Increased duration is consistent with short-term sensitization rather than habituation (42). The cellular mechanism underlying short-term sensitization training in *Aplysia* consist of the activation of serotonergic neurons, which act on the sensory neurons of the reflex circuitry by increasing cAMP intracellular levels (81,82). Hence, increased expression of serotonin and octopamine receptors in the neural circuitry at elevated CO_2_ following habituation could cause a sensitization and produce the increased TWR duration. Furthermore, sensitization-like responses may also be caused by the alteration of metabotropic glutamate receptors. A downregulation of *MGR2* activation, as exhibited in habituated *Aplysia* with OA, could increase production of cAMP in the presynaptic area following habituation training and produce changes in the nervous system consistent with sensitization (83). Trained *Aplysia* exposed to elevated pCO_2_ resulted in specific molecular changes related to the habituation process, for example of *vps-41* coding gene expression. Expression of circular mRNA of *vps-41* improves synaptic plasticity and learning functions by acting as a “sponge” delivering miRNAs to regulate the transcriptome (84,85). Its upregulation under OA and after habituation training could therefore facilitate synaptic changes in the reflex circuitry. Upregulation of *vps-41* could strengthen synaptic connections between sensory neurons and motoneurons through decreased presynaptic transmitter release (86), leading to a sensitization-like behavioural response after elicitation of the reflex. Overall, habituation training while experiencing OA provokes molecular changes that could act on the nervous system in a similar way to that of a sensitization experience in normal conditions, hence modulating the behaviour differently under the influence of changes pH conditions.

*Aplysia* exposed to elevated CO_2_ and administered gabazine did not show restoration of TWR duration consistent with that of control individuals and few changes were observed on the molecular level. Previous studies in molluscs have demonstrated that gabazine could restore impaired behaviours such as burrowing and predator escape (13,87), therefore it is unlikely that the absence of behavioural change is due to the fact that gabazine would not act as a correct antagonist of GABA_A_ receptors. A possible explanation would be that contrary to other behaviours previously investigated in the context of acidification (13,87), the TWR impairment under OA is not caused by interfering with GABA_A_ neurotransmission. This was also seen for another invertebrate which altered behaviour in acidified conditions was not restored with gabazine administration (88). Instead, it is possible that other ligand-gated ion channels (LGICs) performing neurotransmission could have their function altered by the acid-base regulation response in *Aplysia*, such as glutamate or acetylcholine-gated chloride channels which are affected by OA in Aplysia (89). Hence, our results suggest that different mechanisms than changes GABAergic neurotransmission may be responsible for TWR impairment.

Overall, our study reveals links between behavioural alterations and molecular changes occurring in the model neurosystem of *Aplysia* as it is experiencing ocean acidification (OA). Our findings show that the TWR and its modulation through learning is impaired during OA through alterations in neurotransmission between mechanosensory neurons and neurons downstream of the reflex circuitry. Gabazine did not restore TWR duration revealing GABA-ergic neurotransmission not to be the main mechanism responsible for impaired behaviour. By leveraging the simple nervous system of *Aplysia*, we show direct links between near-future predicted ocean conditions and the molecular processes occurring in neurological systems that control for animal behaviour.

## Data availability

The raw sequencing data can be found in BioProject PRJNA1031344. The reviewer link to the data is: https://dataview.ncbi.nlm.nih.gov/object/PRJNA1031344?reviewer=hf09nmkmqj8ebd5b7fa6cu6gbd

## Author contributions

JMS conceived and carried out the experiments with input from CS. JMS analysed the data under the supervision of CS. JMS wrote the paper with input from CS.

## Acknowledgments

JMS and this study were funded by the start-up of CS from the University of Hong Kong. We thank Jay Minuti who provided advice and help to the air mixer system, Sneha Suresh and Arthur Chung who helped with the behavioural experiments, Yan Chit Kam who contributed to the randomization of videos for the blind measurements of TWR duration and Lucrezia Bonzi for her advice on performing molecular analyses. We also thank all the members of the lab for support.

