## Supplementary_Figures for "Ocean acidification alters the transcriptomic response in the nervous system of *Aplysia californica* during reflex behaviour"

| **Name** | **Title** |
| --- | --- |
| Supplementary Figure 1 | Aquarium setup for *Aplysia* housing, composed of a reservoir, experimental tanks and a filtering system; n indicates the range of Aplysia individuals housed per tank; arrows (**>**) indicate the flow direction. The replicate systems were used. |
| Supplementary Figure 2 | pCO_2_ (µatm) in control and elevated CO_2_ (treatment) tanks after completion of the gradual pCO_2_ increase; ambient air was bubbled in the “Control” group tanks whereas a mix of air and CO_2_ (18 cc/min) was bubbled in the “Treatment” group tanks; stars (***) indicate the significant difference between the mean values |
| Supplementary Figure 3 | Tail Withdrawal Reflex (TWR) duration (s) of *Aplysia* in the GABA experiment, reared at elevatwed µatm pCO_2_, as a function of their exposure to gabazine (with) or control water (without); “NS.” stands for “non-significant” and illustrate the absence of a statistical difference between the mean values |
| Supplementary Figure 4 | Mean Tail Withdrawal Reflex (TWR) duration (s) of *Aplysia* in the habituation experiment, reared at control pCO_2_, before (pre-training) and after (post-training) habituation training; “NS.” indicates the absence of significant difference between the mean values |
| Supplementary Figure 5 | Tail Withdrawal Reflex (TWR) duration (s) of *Aplysia* in the learning experiment and before habituation training, as a function of their pCO_2_ conditions (control ~500 µatm or treatment ~1100 µatm); stars (*) indicate the significant difference between the mean values |
| Supplementary Figure 6 | Heatmap correlating expression of genes in clusters produced through the WGCNA analysis and traits such as pCO_2_, GABA exposure, habituation status and mean TWR duration; cell colour is coded by the correlation value (red = positive, blue = negative); upper values are correlation values; lower values in brackets are p-values; bold values indicate significant correlations |

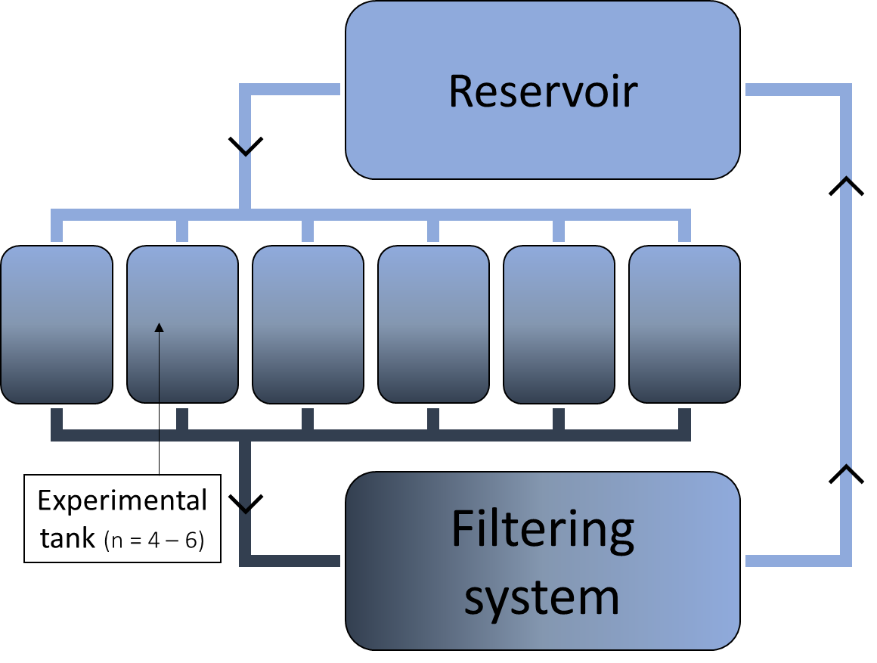
Supplementary Figure 1:

Supplementary Figure 1: Aquarium setup for Aplysia housing, composed of a reservoir, experimental tanks and a filtering system; n indicates the range of Aplysia individuals housed per tank; arrows (**>**) indicate the flow direction. The replicate systems were used.

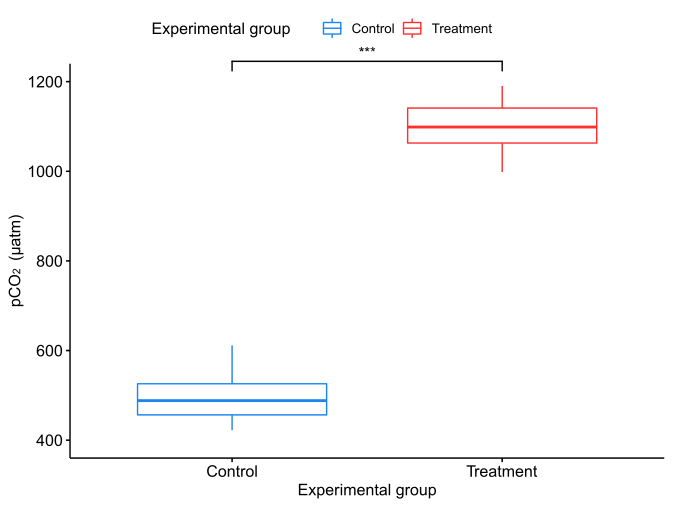
Supplementary Figure 2:

Supplementary Figure 2: pCO_2_ (µatm) in control and elevated CO_2_ (treatment) tanks after completion of the gradual pCO2 increase; ambient air was bubbled in the “Control” group tanks whereas a mix of air and CO2 (18 cc/min) was bubbled in the “Treatment” group tanks; stars (***) indicate the significant difference between the mean values

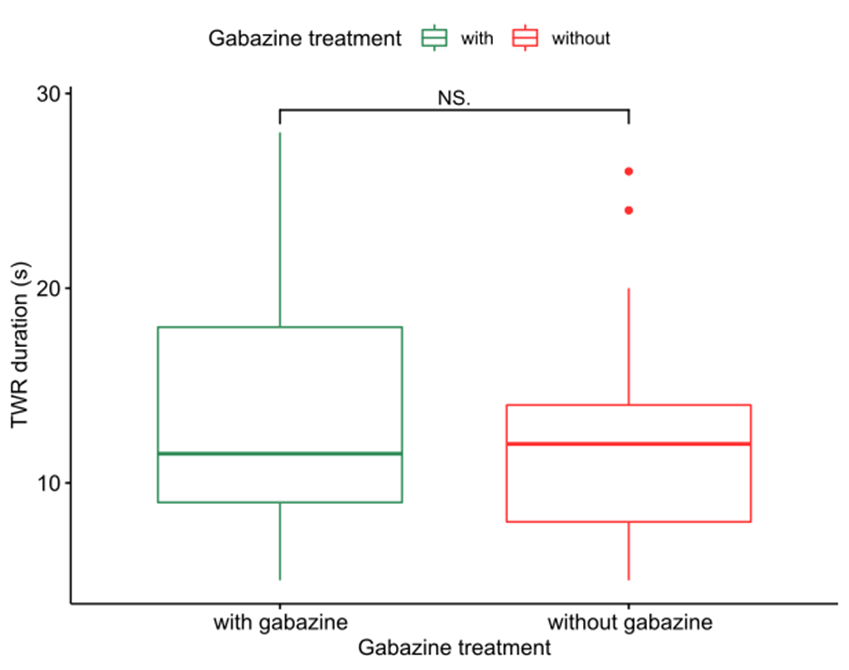
Supplementary Figure 3:

Supplementary Figure 3: Tail Withdrawal Reflex (TWR) duration (s) of Aplysia in the GABA experiment, reared at elevated pCO_2_, as a function of their exposure to gabazine (with) or control water (without); “NS.” stands for “non-significant” and illustrate the absence of a statistical difference between the mean values

Supplementary Figure 4:

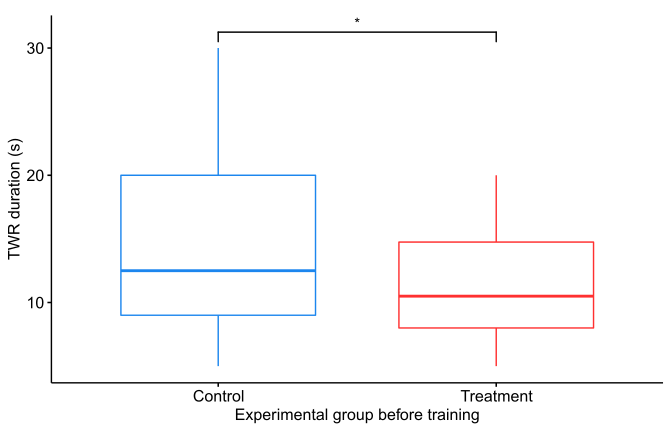

Supplementary Figure 4: Tail Withdrawal Reflex (TWR) duration (s) of Aplysia in the learning experiment and before habituation training, as a function of their pCO_2_ conditions (control ~500 µatm or treatment ~1100 µatm); stars (*) indicate the significant difference between the mean values

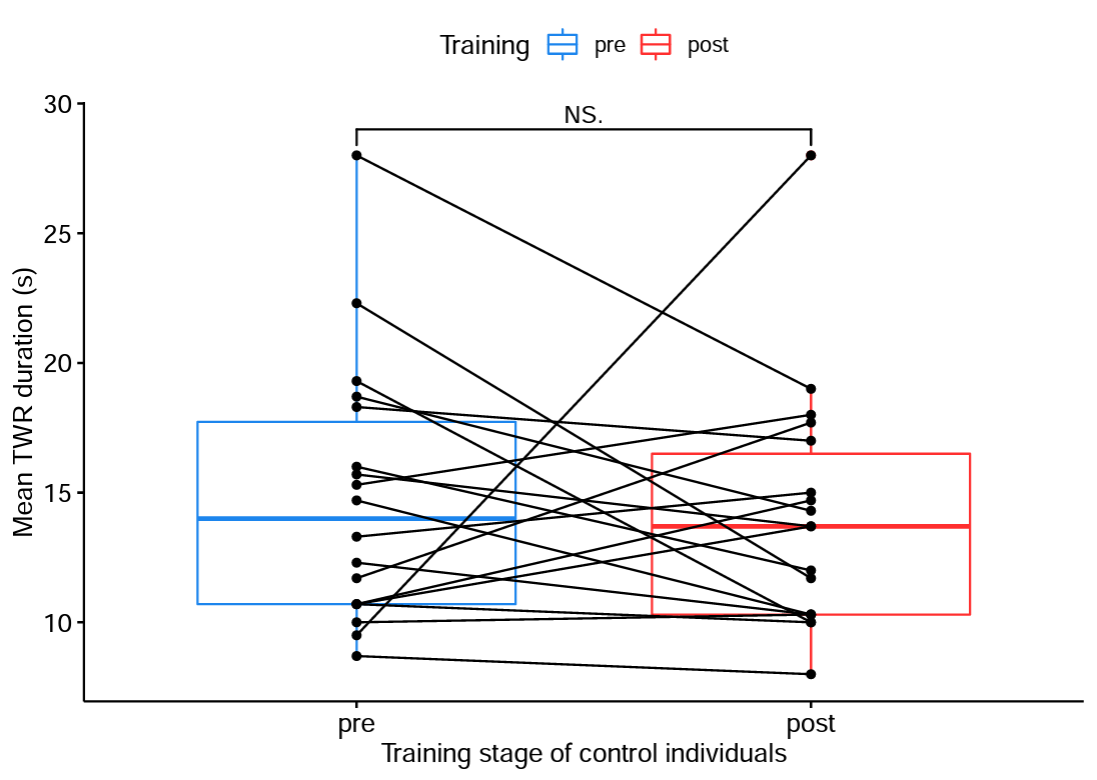
Supplementary Figure 5:

Supplementary Figure 5: Mean Tail Withdrawal Reflex (TWR) duration (s) of Aplysia in the habituation experiment, reared at control pCO_2_, before (pre-training) and after (post-training) habituation training; “NS.”
indicates the absence of significant difference between the mean values

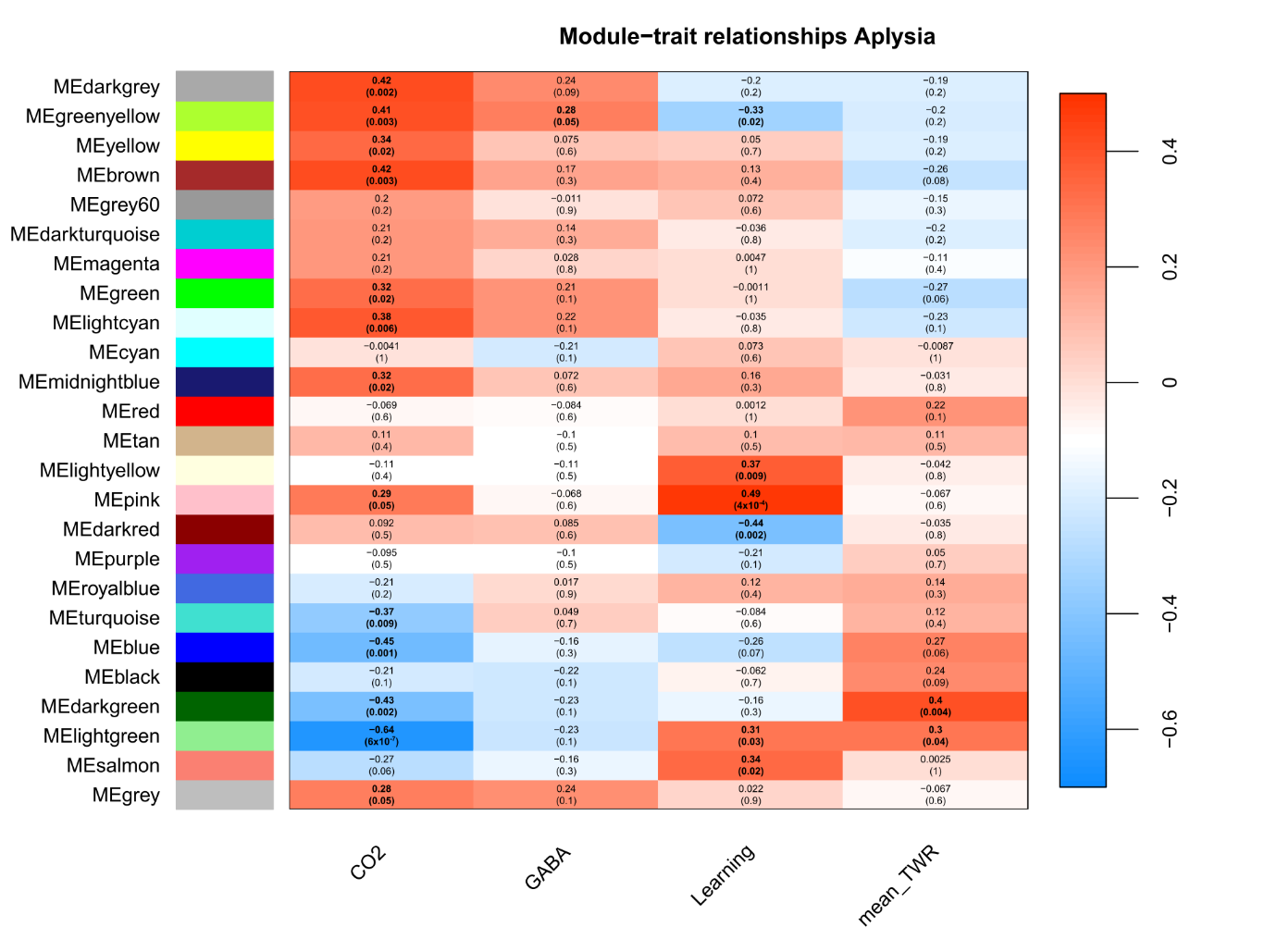
Supplementary Figure 6:

Supplementary Figure 6: Heatmap correlating expression of gene modules by WGCNA analysis and traits such as pCO_2_, GABA exposure, habituation status and individual mean TWR duration. Cell colour is coded by the correlation value (red = positive, blue = negative) based on Pearson correlation. In each cell: upper values are correlation values; lower values in brackets are p-values.
